# Representations of carbon dioxide in the mosquito antennal lobe

**DOI:** 10.1101/2023.03.22.533824

**Authors:** Shefali Goyal, Pranjul Singh, Mudit Gupta, Smith Gupta, Swikriti Saran Singh, Arjit Kant Gupta, Nitin Gupta

## Abstract

Carbon dioxide (CO_2_) is one of the prominent sensory cues used by mosquitoes to find hosts for blood-feeding. CO_2_ is detected on the maxillary palps by capitate peg sensory neurons, whose axons project to the antennal lobe in the brain. Behavioral studies have shown that mosquitoes prefer non-homogenous plumes of CO_2_ over homogenous plumes and CO_2_ greatly enhances the attractiveness of lactic acid, a skin volatile. However, the neural mechanisms underlying these behavioral preferences are not known. Using *in vivo* intracellular recordings from projection neurons and local neurons in the antennal lobe, along with single sensillum recordings from the maxillary palps, we checked the representations of CO_2_ in the first two layers of the *Aedes aegypti* olfactory system. We found that the preference to non-homogeneous plumes of CO_2_ and its synergistic attraction with lactic acid are encoded in the PN population responses. Our results provide a foundation for understanding CO_2_-mediated host-attraction in mosquitoes.

## Introduction

Mosquitoes are vectors for many diseases. For example, *Aedes aegypti* spread Dengue fever (Leta et al., 2018; Messina et al., 2019; Stanaway et al., 2016) and other diseases such as chikungunya, Zika virus, and yellow fever (Bosio et al., 1998; Chompoosri et al., 2016; Hall-Mendelin et al., 2016; Rudnick, 1967). Female mosquitoes spread the disease-causing pathogens while feeding on human blood, which they require to complete their reproductive cycle (Ariani et al., 2015; Dimond et al., 1955; Kogan, 1990). Mosquitoes rely on various sensory stimuli, including visual, thermal, and olfactory cues, and humidity, to find a suitable host for blood-feeding (Cardé, 2015; De Obaldia et al., 2022; M. Z. Liu & Vosshall, 2019; Raji & DeGennaro, 2017; Van Breugel et al., 2015; Zhao et al., 2022). Among these, carbon dioxide (CO_2_) exhaled from the human breath is one of the key stimuli for host-seeking behavior as it activates flight in mosquitoes (Bar-Zeev et al., 1977; Dekker et al., 2001, 2005; Eiras & Jepson, 1991; Geier et al., 1999; Gillies, 1980; McMeniman et al., 2014; P.N. Daykin, 1965). CO_2_ also gates the behavioral attraction to other sensory cues (Acree et al., 1968; Grant & O’ Connell, 1996; McMeniman et al., 2014; Smith et al., 1970; Sorrells et al., 2022).

Odors are detected by olfactory receptor neurons (ORNs) housed in hair-like structures called sensilla (Davis & Sokolove, 1976; Ghaninia et al., 2007, 2008; Kellogg, 1970). Three gustatory receptors – Gr1, Gr2, and Gr3 – are the sole detectors of CO_2_ in *Aedes aegypti* (Jones et al., 2007; McMeniman et al., 2014). These receptors are expressed on cpA neurons present in the capitate peg (cp) sensilla of the maxillary palps (Erdelyan et al., 2012; Grant et al., 1995; Kellogg, 1970). ORN axons send the sensory information to compartments in the antennal lobe (AL) called glomeruli (Couto et al., 2005; Stocker et al., 1990; Vosshall et al., 2000). The estimates of the number of glomeruli in *Aedes aegypti* range between 50 (Ignell et al., 2005) and 80 (Shankar & McMeniman, 2020). In each glomerulus, ORN axons synapse with the dendrites of projection neurons (PNs), which then carry the olfactory information to the mushroom body and the lateral horn (Grabe et al., 2016; Wilson et al., 2004). The axons of cpA neurons innervate a large glomerulus located mediodorsally (MD1) in the AL (Herre et al., 2022; Shankar et al., 2021). Another set of neurons called the local neurons (LNs) facilitate inter-glomerular crosstalk and processing of information within the AL (Bhandawat et al., 2007; Chou et al., 2010; Olsen & Wilson, 2010). Currently there is very limited information about the responses of PNs and LNs to olfactory stimuli in mosquitoes (Lahondère et al., 2020; Melo et al., 2020; Vinauger et al., 2018).

The olfactory cues used by mosquitoes for host-finding include skin emanations such L-lactic acid, 1-octen-3-ol, and acetone (Bernier et al., 1999, 2000; Cork & Park, 1996; Dormont et al., 2013; Zwiebel & Takken, 2004). Interestingly, CO_2_ has a synergistic behavioral effect when given with some of these skin volatiles, especially L-lactic acid (Acree et al., 1968; Eiras & Jepson, 1991; Geier & Boeckh, 1999; Mathew et al., 2013; McMeniman et al., 2014; Smith et al., 1970). Where this synergism arises in the brain remains unclear. CO_2_ is known to be detected only on the maxillary palps, while L-lactic acid responses have been observed in *Ir8a*-expressing neurons on the antennae in mosquitoes (Davis & Sokolove, 1976; Grant et al., 1995; Kellogg, 1970; Raji et al., 2019; Shankar et al., 2021), suggesting that the synergism may be downstream from the sensory neurons.

The preference of an odor can be affected by not only the identity of the odor but also the plume pattern with which it is encountered. Compared to a homogeneous distribution, fluctuating plumes of CO_2_ are more attractive to mosquitoes (Dekker et al., 2001; Geier et al., 1999; Gillies, 1980; Majeed et al., 2017). CO_2_ plume structure is known to affect behavior in other insects also (Barrozo & Lazzari, 2006; Zollner et al., 2004). This makes evolutionary sense as insects encounter non-homogeneous plumes of odor in the natural environment (Murlis et al., 1992). How the stronger response to non-homogeneous plumes is encoded in the mosquito brain is not yet known. These behavioral observations are difficult to explain with labeled-line models of olfactory preferences (Haverkamp et al., 2018), as the same labeled-lines should be activated by the homogenous and the non-homogenous plume patterns.

To gain a comprehensive understanding of the representations of CO_2_ in the mosquito olfactory system, we measured the neural responses of cpA neurons, PNs, and LNs to CO_2_. We compared the responses of these neurons to different temporal patterns of CO_2_. We also measured the responses of AL neurons to CO_2_, L-lactic acid, and their binary blend covering a large fraction of the AL glomeruli. Our results show that the preference to non-homogenous plume patterns and the synergistic responses to the blend of CO_2_ and L-lactic acid are encoded by the PN population. Our results contribute to the understanding of how behavioral preferences to odors are encoded by the mosquito olfactory system and provide a foundation for future work on designing new strategies for controlling mosquitoes.

## Results

### CO_2_ responses of PNs and LNs

We performed *in vivo* whole-cell patch recordings from PNs and LNs in *Aedes aegypti* (P. Singh et al., 2022). In our experimental preparation, we could target cell bodies in medial, dorsal, and dorsolateral regions around the AL. The morphologies of the recorded neurons were detected using dye-fills (see Methods). The glomeruli were identified using the atlas provided by Ignell et al. (Ignell et al., 2005); we use the prefix “I-” before glomerulus names to refer to this atlas. Among more than 200 uniglomerular PNs that we recorded in the dataset, 163 were tested with CO_2_ and had reliable glomerular identifications. Although we could not record from PNs innervating the I-MD1 glomerulus, the PNs in our dataset covered 40 out of the 50 glomeruli reported in the atlas (Ignell et al., 2005). It is possible that the cell bodies of the PNs innervating I-MD1 and the other glomeruli not covered in our dataset are located in the ventral region of the AL, which was not accessible in our experimental preparation.

For each recorded PN, we quantified the average change in the firing rate over the 2s period following the onset of the CO_2_ stimulus (given for 1 s), and compared the responses for all PNs by grouping them according to their glomerular identity **(Figure 1a)**; the average changes in the membrane potentials of these PNs are shown in **Supplementary Figure S1)**. Many types of PNs, such as those innervating I-AM1, I-AC1, I-PD1, I-AD2, and I-AD4 glomeruli, did not respond to CO_2_. I-CD2 and I-PD4 PNs typically showed inhibitory responses to CO_2_ **(Figure 1a, Supplementary Figure S1)**. In 6 glomeruli, for each of which we had at least 8 PNs in our dataset, we checked whether the response magnitudes over all PNs in the glomeruli were statistically different from 0: we found the firing responses of PNs were significantly inhibitory in I-PD6 (mean ± s.d. = 3.34 ± 2.86; P=0.008, n= 8 PNs, signed-rank test), significantly excitatory in I-MD3 (8.26 ± 13.53; P=0.004, n= 16 PNs), and not significantly different from 0 in I-PD2, I-AL2, I-AM2 and I-AL3 glomeruli. However, careful examination of the CO_2_ responses of different PNs within the same glomerulus revealed heterogeneity. For example, among the recordings from two I-PD6 PNs shown in **Figure 1b**, the first PN showed an inhibitory response to CO_2_ and while the second PN showed no response. Similarly, among I-V3 and I-AD5 PNs, some PNs showed excitatory responses while some did not respond to CO_2_ **(Figure 1c, 1d)**. Even among I-MD3 PNs, which were more frequently activated by CO_2_ among the PNs examined in our dataset, there was considerable heterogeneity in the responses **(Figure 1e)**: the first two PNs included in the figure showed clear excitatory responses to CO_2_, the third PN was inhibited by CO_2_, the fourth PN showed a temporally patterned response with an excitatory bout followed by an inhibitory bout, and the fifth PN showed an inhibitory bout followed by a bout of delayed excitation. Given that I-MD3 PNs primarily receive their input from the 1-octen-3-ol-sensitive sensory neurons in the capitate peg sensillum (Singh et al., 2022), it is likely that the excitatory responses to CO_2_ observed in some of the I-MD3 PNs are mediated by lateral excitatory inputs within the AL. In a few PNs, we could test the responses to different concentrations of CO_2_ (0.001, 0.0025, 0.005 and 0.01). The responses were typically stronger for higher concentrations **(Figure 1f)**. This was also true for inhibitory responses (see PN1 and PN2 in **Figure 1f** where CO_2_ 0.01 elicits the strongest inhibitory response).

**Figure 1:**
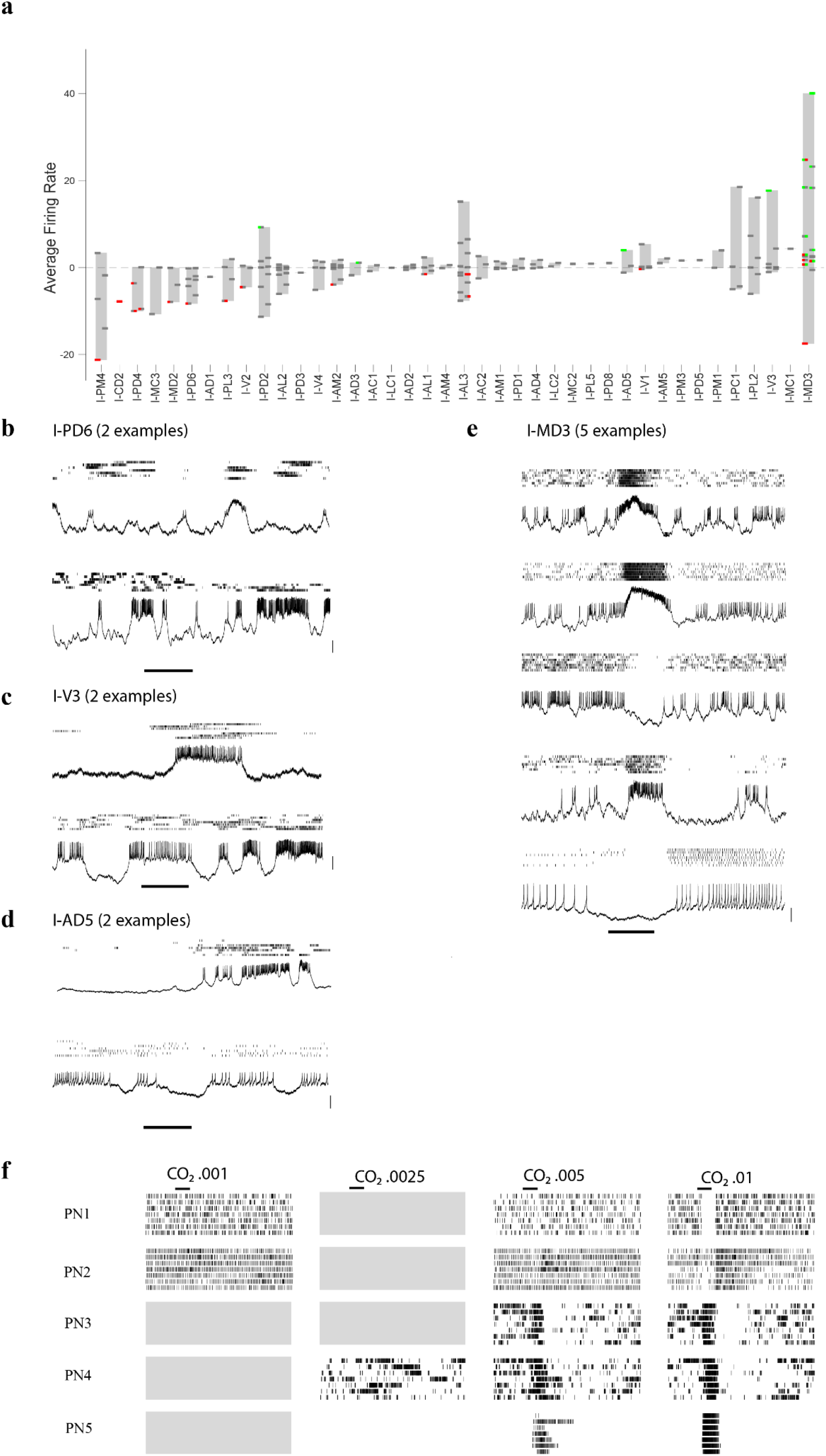
Responses of projection neurons (PNs) to CO_2_. **a** Responses of PNs, organized according to their glomerular identity, to a 1s pulse of CO_2_. Each PN is represented by a thin horizontal rectangle with 2 independently colored halves, corresponding to 1s odor period and 1s post-odor period, respectively. The rectangle is placed at a height corresponding to the average odor-evoked firing rate in the 2s response interval. The color of each half indicates whether the odor-evoked firing in the corresponding period is statistically different from 0 (see Methods); green: significantly >0; red: significantly <0; grey: no statistical difference from 0. **(b-d)** Each figure shows the spike rasters (top) and sample recording traces (bottom) of two different representative PNs innervating I-PD6 (**b**), I-V3 (**c**), and I-AD5 **(d**) glomeruli in response to 1s pulse of CO_2_ (horizontal line). Scale bars, 5 mV. **e** Spike rasters (top) and representative traces (bottom) of 5 PNs innervating I-MD3 glomerulus in response to 1s pulse of CO_2_ (horizontal line). Scale bar, 10 mV. **f** Spike rasters of 5 PNs in response to 1s pulse of CO_2_ (horizontal line) for 4 different concentrations of CO_2_. If a concentration was not tested in a PN, it is indicated by an empty grey region.

We expected that the lateral inputs underlying the excitatory and inhibitory responses to CO_2_ in I-MD3 and other PNs, which do not receive direct input from the CO_2_-sensitive sensory neurons on the maxillary palps, could come from LNs. Our patch-recordings in the AL also included LNs, whose identity was confirmed by dye fills. Overall, our dataset included 49 LNs that were tested with CO_2_. Consistent with our expectation, we found that some LNs showed excitatory responses, some showed inhibitory responses, and some did not respond to CO_2_ **(Figure 2a, b)**. The frequency of excitatory responses in LNs was slightly above 20% (10 out of 49 PNs), which was more than that seen in PNs (14 out of 163 PNs, i.e., less than 10%). A similar proportion of LNs (9 out of 49) was also inhibited by CO_2_. Overall, these results indicate that CO_2_ activates or inhibits a sizeable fraction of LNs, which could then spread the activity to diverse PNs in the AL.

**Figure 2:**
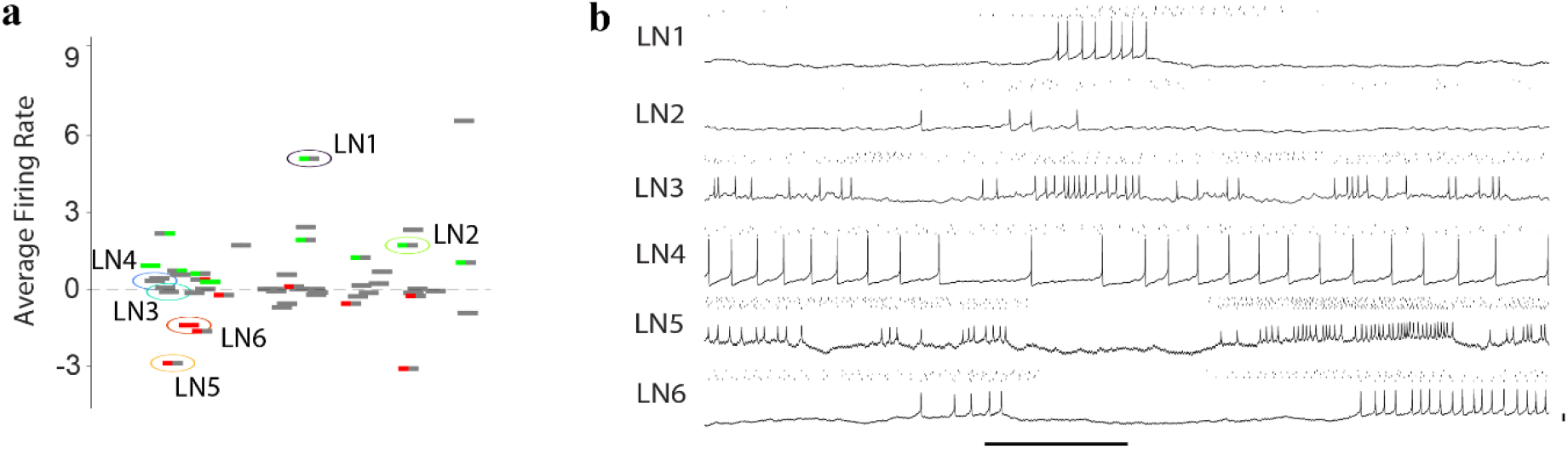
Responses of local neurons (LNs) to CO_2_. **a** Responses of 49 LNs to a 1s pulse of CO_2_, plotted using the same representation as used in Figure 1a. Six representative LNs are encircled and labeled LN1-6. **b** Spike rasters and recording traces of these six LNs in response to 1s pulse of CO_2_ (horizontal line). Scale bar, 10 mV.

To further explore this idea, we measured the responses to odorants cyclopentanone and 3-methyl cyclopentanone, which are known to mimic CO_2_ responses in the maxillary palps (Tauxe et al., 2013). We made single-sensillum recordings from the capitate peg sensillum on the maxillary palps and measured the responses of the CO_2_-sensitive cpA neurons to these odorants (see Methods). In agreement with previous results, we found that the responses of cpA to cyclopentanone or 3-methyl cyclopentanone were similar to the responses to CO_2_ **(Supplementary Figure S2a, b)**. Although CO_2_ is not detected on the antennae, the other odorants may be detected by additional sensory neurons on the antennae. PN recordings from two mosquitoes in which the maxillary palps were blocked from odor stimulation (see Methods) showed depolarization of the membrane potential in response to cyclopentanone but not in response to CO_2_, suggesting that cyclopentanone activates some sensory neurons on the antennae **(Figure 3a)**. The axons of the sensory neurons from the palps and from the antennae project to different glomeruli in the antennal lobe (Herre et al., 2022; Ignell et al., 2005). However, LNs can provide cross-talk across these glomeruli. **(Figure 3b)** shows odor responses of LNs in intact mosquitoes in which CO_2_ and cyclopentanone were tested. Some of these LNs responded differently to the two odorants: LN2 showed an excitatory response to CO_2_ and LN5 showed an inhibitory response to CO_2_ but neither responded to cyclopentanone. On the other hand, LN4 and LN8 showed stronger responses to cyclopentanone than to CO_2_. Similarly, some differences were noticeable in the LN responses to CO_2_ and 3-methyl cyclopentanone **(Supplementary Figure S3a)**. Next, we examined the responses of PNs to these odorants, and consistent with the idea of lateral interactions mediated by LNs, we found diverse PN responses to these odorants **(Figure 3c, Supplementary Figure S3b)**. For instance, I-AD2 and I-V3 PNs did not respond to CO_2_ but showed excitatory responses to cyclopentanone and 3-methyl cyclopentanone; some PNs, such as those belonging to I-PC1 and I-MD3 glomeruli, responded to these odors with different temporal patterns.

**Figure 3:**
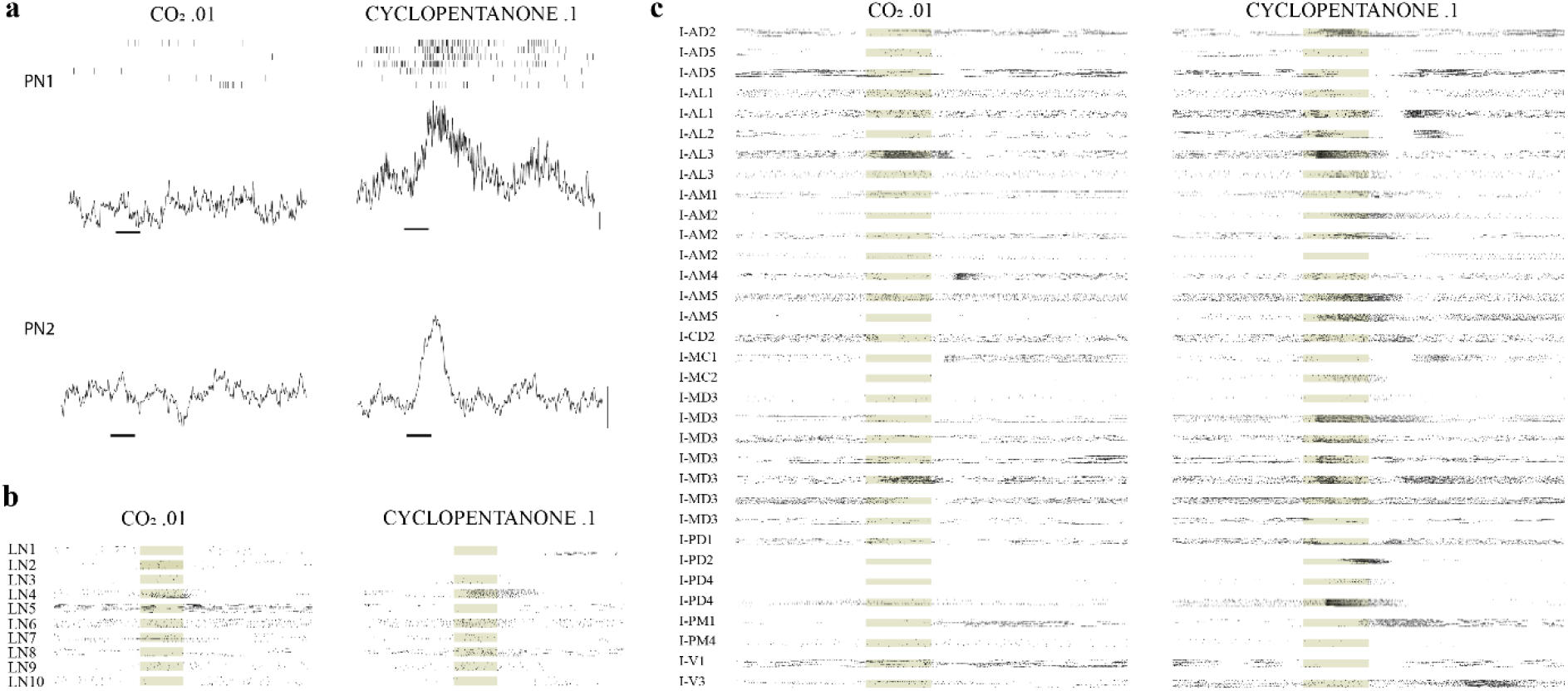
Comparing responses to CO_2_ and cyclopentanone. **a** Responses of PNs to CO_2_ and cyclopentanone in two mosquitoes in which the antennae but not the maxillary palps were stimulated with the odors. Spike rasters and the trial-averaged membrane potential traces (after 20 Hz low-pass filtering) are shown. No spikes were observed in PN2. Note the odor-evoked depolarization in the membrane potential in response to cyclopentanone, but not CO_2_, in both neurons. Odor pulses were 1s long (horizontal line). Scale bar, 1 mV. **b** Spike rasters showing responses of LNs to 1s pulses of CO_2_ and cyclopentanone (shaded background). **c** Spike rasters showing responses of PNs, sorted by the glomerular identity, to 1s pulses of CO_2_ and cyclopentanone.

### Neural responses to temporally patterned CO_2_ stimuli

We asked if the higher behavioral attraction to non-homogenous plumes of CO_2_ can be explained by the neural responses in the early stages of olfactory processing. We performed experiments with three temporal patterns of CO_2_ stimuli: a single pulse of 4s duration, four pulses of 1s duration (separated by 1s gaps), and eight pulses of 0.5s duration (separated by 0.5s gaps). The first of these patterns is the most homogenous while the last one is the most heterogenous; note that the amount of odor delivered is equal in all three patterns (4s each). Using single-sensillum recordings (SSRs), we checked the spiking responses of cpA neurons on the maxillary palps to the three patterns **(Figure 4a)**. Combining data from 6 SSRs **(Figure 4b)**, we observed that the total number of spikes generated by the three patterns were not different from each other (single vs four pulses: P=0.84; single vs eight pulses: P=0.44; four vs eight pulses: P=0.16; n=6 in each comparison; signed-rank tests). This result confirms that the response strength of the peripheral neurons does not change significantly with the temporal pattern of the CO_2_ stimulus if the total amount of stimulus remains constant.

**Figure 4:**
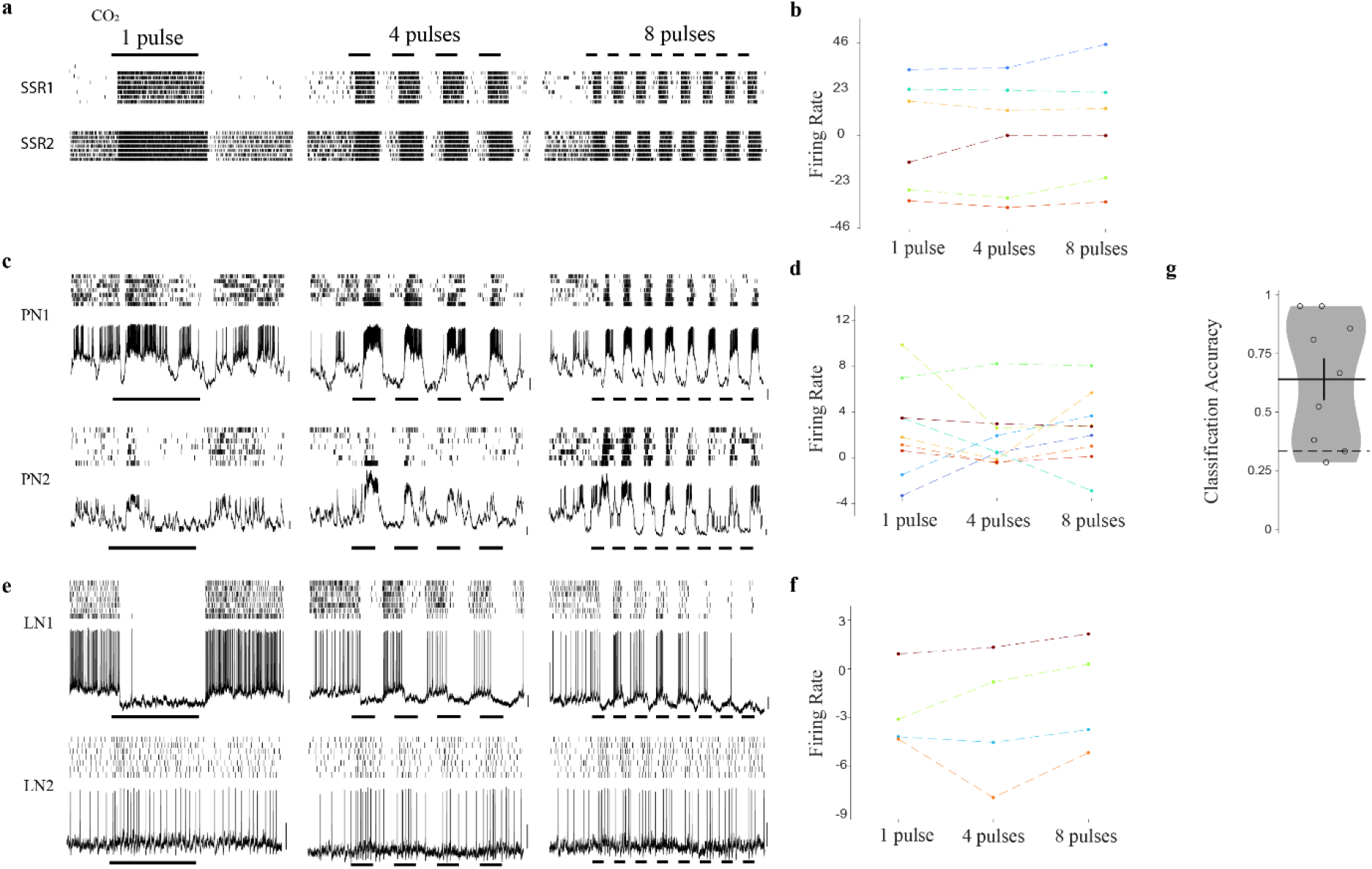
Responses of sensory neurons, PNs, and LNs to temporally patterned CO_2_ stimuli. **a** Spiking responses of cpA sensory neurons from two representative single sensillum recordings in the maxillary palps for 3 different temporally patterns of CO_2_ stimulation: 1 pulse of 4s duration, 4 pulses (middle) of 1s duration each, and 8 pulses of 0.5s each (horizontal lines). **b** Average changes in the firing rates of cpA sensory neurons (n = 6) during the 8s response interval (compared to the background) in response to the 3 temporal patterns of CO_2_ stimuli. **c, e** Spikes rasters and recording traces from two representative PNs (**c**) and two representative LNs (**e**) in response to the 3 temporal patterns of CO_2_ stimuli. **d, f** Average changes in the firing rates of PNs (n = 9, **d**) and LNs (n = 4, **f**) during the 8s response interval (compared to the background) for the 3 stimuli. **g** Prediction accuracy of each PN (n = 9) for the 3 temporally patterned stimuli of CO_2_. Dashed line shows the chance level (1/3).

Next, we asked if the neural responses at the next layer of the olfactory processing are sensitive to the plume structure of CO_2_. We tested the responses of PNs to the same three patterns of CO_2_ as described above. The responses of the PNs appeared to follow the temporal pattern of the stimulus **(Figure 4c)**. However, over the set of 9 CO_2_-responsive PNs in which this experiment could be performed **(Figure 4d)**, the overall strength of the spiking response was not different for the three patterns (single vs four pulses: P=0.65; single vs eight pulses: P=1.0; four vs eight pulses: P=0.25; n=9 in each comparison; signed-rank tests). We also checked the responses of LNs **(Figure 4e)** and did not observe a reliable difference in the total spiking response for the three patterns (single vs four pulses: P=1.0; single vs eight pulses: P=0.38; four vs eight pulses: P=0.13; n=4 in each comparison; signed-rank tests; **Figure 4f**).

Thus, even at the antennal lobe level, the magnitudes of the neural responses do not appear to change with the temporal patterns of CO_2_ stimulation. However, the information about the temporal pattern of the stimulus appeared to be retained in the temporally patterned spikes of PNs, which could carry this information to downstream neurons. We quantified this information by asking if the stimulus pattern of a recording could be predicted by looking at the spiking responses in each PN (see Methods).

Compared to the chance level of 0.33 in predicting one of the three stimulus patterns, we could predict the stimulus patterns of recordings with a significantly higher accuracy of 0.64 ± 0.27 (P = 0.02, n = 9 PNs, signed-rank test; **Figure 4g**). Thus, PN response patterns indeed contain information about the temporal patterns of stimuli, which can potentially be used by the downstream neurons in the higher olfactory centers to generate stronger behavioral preference to non-homogenous patterns of CO_2_.

### Sensory responses to CO_2_, L-lactic acid, and their mixture

We next sought to understand if the synergistic behavioral responses previously observed for the mixture of CO_2_ and L-lactic acid originate in the first two stages of the olfactory processing. Because CO_2_ does not activate sensory neurons on the antennae but activates cpA sensory neurons on the maxillary palps, we examined the responses of maxillary palp neurons for possible synergism with L-lactic acid. Using single-sensillum recordings from the capitate peg sensilla to detect cpA, cpB and cpC spikes (see Methods), we checked the responses of the three neurons to CO_2_, L-lactic acid, and their mixture **(Figure 5a)**. We also tested the responses to water as a control for L-lactic acid (which was dissolved in water).

**Figure 5:**
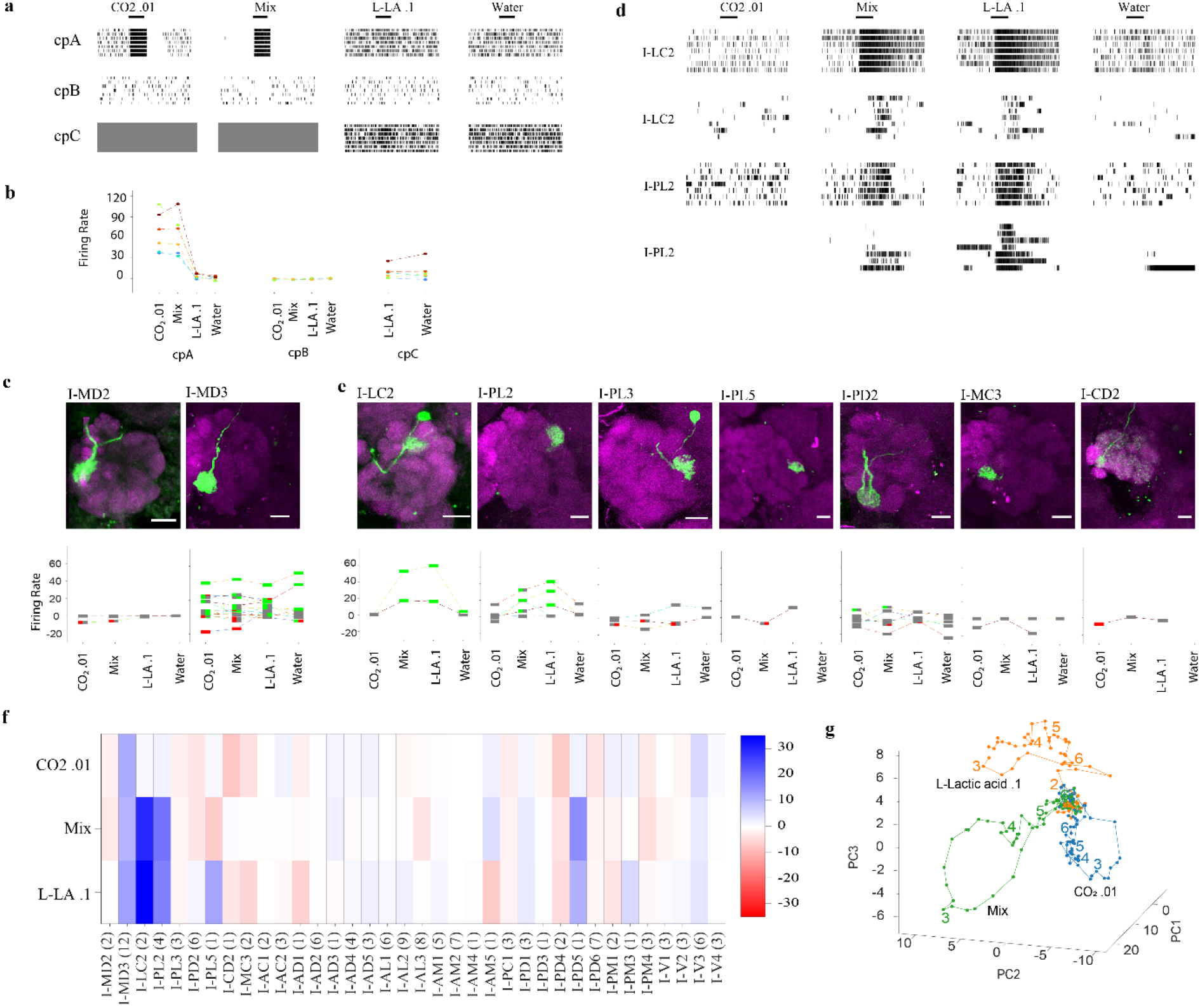
Responses of sensory neurons, PNs, and LNs to CO_2_, L-lactic acid, and their mixture. **a** Spiking responses of cpA, cpB and cpC sensory neurons from a representative single sensillum recording to 1s pulses of CO_2_, L-lactic acid (L-LA), their mixture (mix), and water (the solvent control for L-lactic acid). Responses of cpC to CO_2_ and the mixture could not be assessed (grey boxes). **b** Odor-evoked changes in the firing rates of cpA, cpB and cpC neurons from 6 different sensilla for the same set of odors. **c** Top, representative histological images showing the glomerular innervation of I-MD2 and I-MD3 PNs. Each image is a maximum intensity projection of z-sections covering the antennal lobe. The neuropil labeled using anti-N-cadherin antibody is shown in magenta, and the dye-filled PN is shown in green. Scale bar, 20 µm. Bottom, odor-evoked changes in the firing rates of I-MD2 and I-MD3 PNs. **d** Spiking responses of two representative I-PL2 and two I-LC2 neurons to the same set of odors. **e** Histological images and odor-evoked changes in the spiking rates of PNs innervating 7 different glomeruli that receive direct input from *Ir8a*-expressing sensory neurons. Scale bar, 20 µm. **f** Heat map summarizes the average change in the firing rates of 36 types of PNs to CO_2_, L-lactic acid, their mixture. The number of PNs in each glomerulus tested with the three odors is mentioned in parenthesis. **g** Trajectories of PN population responses to CO_2_ (blue), L-lactic acid (orange), and their mixture (green) in the three-dimensional principal component space. Trajectories are plotted for the 0-6s interval from 10s trials; the odor was given during the 2-3s interval. All trajectories hover around the origin during the spontaneous activity before the odor pulse, but start to diverge after the odor onset at the 2s time-point, and return to the origin after a few seconds. Numbers in the graph represent the time-points (some numbers are hidden behind the dense region near the origin).

We found that cpA neurons did not respond to L-lactic acid **(Figure 5b)**; they responded to the mixture of CO_2_ and L-lactic acid, but the response was not different from the response to CO_2_ alone (P=0.69, n= 6 sensilla, signed-rank test). We did not observe any response in cpB to CO_2_, L-lactic acid, or their mixture **(Figure 5b)**. In case of cpC neurons, it is difficult to identify their spikes (which are very small in amplitude) when cpA neurons are firing at high rates, as happens in the presence of CO_2_. Therefore, we did not analyze cpC spikes for CO_2_ and the mixture stimuli, but it has been shown previously using experiments in Gr3-mutant mosquitoes (which lack cpA spikes) that cpC neurons do not respond to CO_2_ (McMeniman et al., 2014; Singh et al., 2022). We checked the responses of cpC neurons to L-lactic acid and water **(Figure 5b)** and found no or weak responses that were not different across the two stimuli (P=0.56, n= 6 sensilla, signed-rank test), which suggests that cpC neurons do not respond to L-lactic acid per se but respond weakly to water. Considering that cpC neurons do not respond to either CO_2_ or L-lactic acid, they are unlikely to be a site for generating a synergic response to the mixture. Overall, we did not find any evidence of synergism in the three types of neurons on the maxillary palps. These results suggest that the synergism between CO_2_ and L-lactic acid is not encoded at the periphery and perhaps originates downstream of the sensory neurons.

### PN responses to CO_2_, L-lactic acid, and their mixture

At the next layer, we examined the responses of PNs in the AL. We first considered if the three glomeruli receiving direct input from the maxillary palps are involved in the synergism. While we could not record from I-MD1 PNs as mentioned earlier, we were able to check the responses of I-MD2 and I-MD3 PNs. We had two I-MD2 PNs in which the responses to CO_2_, L-lactic acid, and the mixture were recorded **(Figure 5c)**. These PNs did not show excitatory responses to any of these odors (one of them showed inhibition to CO_2_ and the mixture). I-MD3 PNs, which receive input from cpC neurons (Singh et al., 2022), responded equally to L-lactic acid and water (P=0.89, n= 9 PNs, signed-rank test; **Figure 5c**), suggesting that these PNs, like cpC neurons on the maxillary palps, may also be responding to water present as the solvent for L-lactic acid and not to L-lactic acid per se. As discussed above, I-MD3 PNs showed a variety of responses to CO_2_ (**Figure 1e**); overall, these responses were similar in magnitude to the responses to the mixture of CO_2_ and L-lactic acid (P=0.625, n= 14 PNs, signed-rank test; **Figure 5c**). Thus, we did not find any evidence of synergism in I-MD2 and I-MD3 PNs.

We next considered the PNs that receive input from *Ir8*a-expressing sensory neurons on the antenna, as Ir8a is required for the detection of L-lactic acid. Recent studies have shown that *Ir8a*-expressing sensory neurons innervate a subset of glomeruli in the AL (Herre et al., 2022; Shankar et al., 2021; Shankar & McMeniman, 2020), which map onto 11 glomeruli (I-LC1, I-LC2, I-PL1, I-PL2, I-PL3, I-PL4, I-PL5, I-PD2, I-MC3, I-CD2, I-CD4) in the brain atlas we used (Ignell et al., 2005). We were able to record the responses of CO_2_, L-lactic acid, and the mixture from PNs belonging to 7 of these 11 glomeruli (I-LC2, I-PL2, I-PL3, I-PL5, I-PD2, I-MC3, I-CD2). We found that I-PL2 and I-LC2 PNs showed excitatory responses to L-lactic acid (these responses were stronger than their responses to water; **Figure 5d**). But these cells did not respond to CO_2_ and their responses to the mixture were similar to the responses to L-lactic acid **(Figure 5e)**. I-PL3, I-PL5, I-MC3, and I-PD2 PNs did not show clear responses to L-lactic acid, CO_2_, or the mixture (**Figure 5e**). I-CD2 PN showed an inhibitory response to CO_2_ but no response to L-lactic acid or the mixture **(Figure 5e)**. We were also able to test the mixture responses of one biglomerular PN and one multiglomerular PN that innervated some of the glomeruli receiving Ir8a input, but in these neurons also the mixture responses were not stronger than the responses to CO_2_ or L-lactic acid **(Supplementary Figure S4)**. Thus, although some of the glomeruli receiving input from Ir8a-positive sensory neurons responded to L-lactic acid, we did not detect strong synergism with CO_2_ in the 7 glomeruli that we could test.

Lastly, we checked the PN responses in glomeruli that receive input from other sensory neurons on the antenna. The average responses of these PNs to CO_2_, L-lactic acid, and the mixture are shown in **Figure 5f**. In each glomerulus, the average PN response to the mixture was similar to or weaker than the responses to the two components. Overall, among the 36 different glomeruli in which we could test PN responses, we did not find obvious synergism in the mixture responses of the individual PNs.

### The mixture is encoded differently in the PN population

Although individual PNs seemed unable to distinguish the CO2-lactic acid mixture from the individual components, it is possible that the PN population as a whole is able to represent the mixture differently. We visualized the population responses to the mixture and the individual components by plotting the response trajectories over time in a high-dimensional space, with each dimension corresponding to one PN (but reduced to 3 dimensions using the principal component analysis; see Methods). The response trajectories for the odors start at the same origin point, corresponding to the spontaneous activity of the PN population, but after encountering the 1 s-long odor pulse starting at the 2s timepoint, they diverge in different directions to return to the origin around the 5-6s timepoint. The response to the mixture (green) traced a completely different trajectory compared to the trajectories for CO_2_ and L-lactic acid **(Figure 5g)**, suggesting that PN population is able to represent the mixture differently from the individual components.

To quantify how well a neuron or a population of neurons can differentiate the mixture from the two components, we used an odor-classification analysis. We checked if the mixture response in each trial was closer to the template created for the mixture from the remaining trials compared to the templates for the individual components (see Methods). As the mixture response in each trial is compared against 3 templates, the chance level is 1/3. When we performed this analysis for each PN separately, we found that the average accuracy over the 7 trials was 0.36 ± 0.31 (mean ± s.d., n = 151 PNs), which was not significantly different from the chance level of 1/3 (P = 0.27, n = 151 PNs, signed-rank test). This result quantitatively supports our earlier observation that individual PNs in general are not good at differentiating the mixture from the individual components. However, when we considered the responses of all PNs together, as a population, we observed perfect classification with the accuracy value of 1 in each of the 7 trials (P = 0.016, n = 7; signed-rank test against 1/3). Thus, the PN population accurately differentiates the CO_2_-lactic acid mixture from the individual components.

## Discussion

Synergistic responses to mixtures of different odorants have been observed in a variety of insects (Liu et al., 2019; Piñero et al., 2008, 2020; Varela et al., 2011). In mosquitoes, the best-known example is the synergism between CO_2_ and L-lactic acid (Acree et al., 1968; Eiras & Jepson, 1991; Geier & Boeckh, 1999; Mathew et al., 2013; Raji et al., 2019; Smith et al., 1970). Although it has been known from morphological studies that the axons from the antennal sensilla (that respond to L-lactic acid) and from the palp sensilla (that respond to CO_2_) project to different glomeruli in the antennal lobe (Anton et al., 2003; Ignell et al., 2005), it was not clear where the synergistic response to the mixture arises. We did not find notable synergism at the level of individual PNs or LNs. Although the mixture response was similar to one of the component responses in most neurons, these responses were not always identical, leaving open the possibility that small differences at the level of individual neurons can result in a large difference at the population level. Indeed, when we looked at the dynamic responses of the whole PN population, we saw a clear divergence between the response to the mixture and responses to the components (**Figure 5g**). Consistent with this divergence, we found that the PN population responses could be used to accurately classify the odor mixture. Our results extend the idea of a spatiotemporal code for odors in the insect antennal lobe (Riffell et al., 2009; Saha et al., 2013; Wilson, 2005) to mosquitoes.

The PN responses are read out by downstream neurons in the lateral horn and the mushroom body, where the spatiotemporal patterns in the PN population can result in the activation of different sets of downstream neurons (Kymre et al., 2021; Perez-Orive et al., 2002; Strutz et al., 2014). These differences in the PN population may be enough to drive a different motor program for the mixture compared to the components (Jung et al., 2015). In our experiments, the two odors were given at the same time. As the olfactory system can make use of the fine differences in the arrival times of different odorants within a mixture (Sehdev et al., 2019), it would be interesting to check in future work if this temporal aspect is also important in the synergistic responses to CO_2_ and L-lactic acid.

Odor separation by the PN population is not directly inherited from sensory neurons but rather involves lateral processing in the antennal lobe (Bhandawat et al., 2007). This processing is enabled by a diverse population of LNs present in the area (Chou et al., 2010), involving both excitatory and inhibitory connections (Das et al., 2017; Mohamed et al., 2019). Lateral inhibitory networks are also involved in odor discrimination in the olfactory bulb in vertebrates (Abraham et al., 2010; Kay et al., 2009). In our analysis, more than 20% of the recorded LNs showed excitatory responses to CO_2_ and another ∼20% showed inhibitory responses, suggesting a widespread involvement of LNs in shaping the CO_2_ dynamics in the mosquito antennal lobe.

Shankar et al. have recently examined the responses of *Ir8a*-expressing olfactory sensory neurons using CaMPARI2, focusing on the axon processes of these sensory neurons in 12 different glomeruli in the antennal lobe (Shankar et al., 2021). They detected responses to the mixture of L-lactic acid and CO_2_ in some of these glomeruli, including PL6 and PL4, which correspond to I-LC2 and I-PL2, respectively, in our reference atlas (Ignell et al., 2005; Shankar & McMeniman, 2020). Although the *Ir8a*-expressing neurons were targeted because they are known to be required for the detection of L-lactic acid (Raji et al., 2019), the authors surprisingly did not find responses to L-lactic acid in any of these neurons (Shankar et al., 2021). Because their experiments were performed using CaMPARI2, which required permanent photoconversion after odor exposure and post-hoc imaging, the responses of the sensory neurons to L-lactic acid and the mixture were measured in different mosquitoes. In contrast, by using real-time measurements of the odor responses with the help of in vivo electrophysiology, we could measure the PN responses to multiple odors in the same mosquito. While we found responses to the CO_2_-L-lactic acid mixture in I-PL2 and I-LC2 PNs, we also found responses to L-lactic acid in the same PNs. The simplest explanation of our results is that L-lactic acid alone can activate some of the *Ir8a*-expressing sensory neurons, which in turn activate the PNs in the corresponding glomeruli. Future studies measuring the activity of Ir8a-positive sensory neurons with electrophysiology or live calcium/voltage imaging will help in verifying their responses to L-lactic acid.

One shortcoming of our experimental dataset is that despite having more than 200 PNs belonging to 40 different glomeruli, it did not contain PNs innervating I-MD1 glomerulus, which is known to receive input from cpA neurons (Herre et al., 2022; Ignell et al., 2005; Shankar et al., 2021). This was surprising (and disappointing) to us, especially considering that I-MD1 is the largest glomerulus in the antennal lobe (Ignell et al., 2005; Shankar & McMeniman, 2020). In contrast, PNs innervating I-MD2 and I-MD3 glomeruli, which also receive input from the maxillary palps and are considerably smaller in size than I-MD1, were observed 3 and 16 times, respectively. Therefore, it seems unlikely that the lack of I-MD1 is related to incomplete sampling. Rather, we think the reason may be related to a different location of the cell bodies of I-MD1 PNs compared to I-MD2 and I-MD3 PNs. In our experimental preparation, to allow access to the recording electrode, the cuticle was removed above the dorsal side of the brain, while avoiding damage to the antennal nerve located towards the anterior-ventral side. This preparation allowed us to target cell bodies in the dorsal and lateral regions of the antennal lobe. Based on the outcome of our experiments, it appears likely that the cell bodies of I-MD1 PNs are located more ventrally and hence were not accessible.

Nevertheless, we detected responses to CO_2_ in some of the other types of PNs. In *Drosophila*, even though the sensory neurons expressing the CO_2_-sensitive gustatory receptors project to a single glomerulus, called the V glomerulus (Kwon et al., 2007; Suh et al., 2004), recent work has shown that CO_2_ also activates other glomeruli in the antennal lobe in addition to the V glomerulus (Zocchi et al., 2022). Interestingly, these additional glomeruli include some glomeruli (VA2 and DM1) that also receive input from other sensory neurons housed in the same sensillum (ab1) that houses the CO_2_ sensory neurons. This is in line with our observations in mosquitoes, as we also found CO_2_ responses in I-MD3 glomerulus, which receives input from cpC neurons located in the same sensillum as cpA neurons (Singh et al., 2022). In *Drosophila*, the CO_2_ responses in VA2 and DM1 glomeruli are expected to originate from lateral interactions among the sensory neurons (Zocchi et al., 2022), which may also be the case in mosquitoes. We also observed inhibitory responses to CO_2_ in many glomeruli, which points to lateral inhibition, possibly mediated by the GABAergic LNs.

We found heterogeneity in the CO_2_ responses of PNs innervating the same glomerulus (**Figures 1b-e**). This points to functional diversity among sister PNs within a glomerulus (Dhawale et al., 2010). Although sisters PNs in many glomeruli in *Drosophila* appear to be very similar (Kazama & Wilson, 2009), remarkable functional diversity has been reported in a CO_2_-sensitive glomerulus: there are at least three different types of PNs innervating the V glomerulus, which differ in their morphologies, neurotransmitters, and responses to CO_2_ (Lin et al., 2013). In the moth *Helicoverpa armigera*, sister PNs innervating the CO_2_-sensitive labial pit organ glomerulus (LPOG) have also been shown to have morphological differences (Chu et al., 2020). Thus, functional diversity among CO_2_-sensitive PNs appears to be a general feature of the insect brain.

## Methods

### Animal stocks

All experiments were conducted with wild-type *Aedes aegypti* (Liverpool strain) mosquitoes. The larvae were fed with powdered fish food (Tetrabits). Mosquitoes used for experiments were females aged 4-9 days post-eclosion, maintained at a temperature of 25±5°C and relative humidity of 60±15%. They were fed with 10% sucrose solution and blood-fed using adult wild-type mice *Mus musculus BALB/c*.

### Animal preparation for *in vivo* whole-cell patch

Mosquitoes were anesthetized on ice for about a minute and mounted on a small plastic dish with a square hole. Wings were fixed on both sides of the plastic dish and legs were pushed down through the hole and glued with wax. The head of the mosquito was raised to the level of the thorax and fixed in this orientation with epoxy resin. A thin plastic sheet with a trapezium-shaped hole was placed over the head such that it covered the entire body except for the head (the region between the Johnston’s organs and the thorax). The gaps between the mosquito and the plastic sheet were sealed with epoxy resin. The plastic sheet acted as a separator, such that the antennal lobes could be accessed above the sheet after removing the cuticle and perfusing saline, and the sensory organs (antennae and palps) could be kept dry for odor stimulation below the sheet. The mosquito was then fixed on a recording chamber with wax such that the head was accessible for dissection. A small window was cut in the head just below the Johnston’s organs to access the antennal lobe neurons for *in vivo* whole-cell patch recordings. After the surgery, the head was continuously perfused with a saline solution at a flow rate of 1-2 ml/min. The composition of the saline was as follows: 12.5 mM N-tris(hydroxymethyl)methyl-2-aminoethane-sulfonic acid (TES), 8 mM glucose, 2 mM sucrose, 120.5 mM sodium chloride, 6.25 mM trehalose dihydrate, 17 mM sodium bicarbonate, 0.6 mM sodium dihydrogen phosphate monohydrate, 3.15 mM potassium chloride, 1.6 mM calcium chloride dihydrate, 2.9 mM magnesium chloride hexahydrate. The osmolarity was adjusted to 300-305 mOsm and pH was finally adjusted to 7.3 after bubbling with a 95% O_2_/5% CO_2_ stream.

In some experiments, the odor stimulation of the maxillary palps was blocked by either severing them with a sharp blade or covering them with epoxy resin during the experimental preparation. This manipulation removed the olfactory activity of the maxillary palps while leaving the antennal activity intact.

### Electrophysiological recordings from PNs and LNs

For *in vivo* whole-cell patch recordings, an amplifier (Multiclamp, Axon) and data acquisition system (Digidata 1550A, Axon) were used. The animal was mounted on a fixed stage under a Nikon FN1 microscope with a 40X water objective. Images were obtained with a Nikon DS-Qi2 camera. Patch electrodes were pulled with an electrode puller (Sutter P-1000) and had a resistance of about 6-9 MΩ. They were filled with internal patch solution including 140 mM potassium aspartate, 10 mM HEPES buffer, 4 mM MgATP, 0.5 mM Na_3_GTP, 1.1 mM EGTA, 10 mM potassium hydroxide, 1 mM potassium chloride. The solution also included 0.5 % biocytin or 0.1 % Lucifer yellow for staining the neurons. The osmolarity was adjusted to approximately 295 mOsm and pH was adjusted to 7.3.

### Odor stimulation

Odors were delivered to the mosquito through polytetrafluoroethylene (PTFE) tubing at a distance of about 8mm from the antennae at a final flow rate of 2 L/min. This included a continuous 1.8 L/min stream of activated charcoal-filtered clean air and 0.2 L/min of a variable stream that switched between odorized air or clean air during the experiments. The switching was performed using a 3-way solenoid valve (Parker Hannifin), which was controlled by the Digidata 1550A data acquisition system (Molecular Devices). CO_2_ was delivered from a cylinder. Other odorants including L-lactic acid (Sigma, 27714), cyclopentanone (Merck, 802670), 3-methyl cyclopentanone (Sigma, 157643) were placed in 50 ml glass bottles. The odor delivery system included a fixed air dilution of 0.1 because of the mixing of the odorized stream (0.2 L/min) into the constant background stream (1.8 L/min). Further dilutions, if required, were performed by diluting (volume/volume) the odorant in the glass bottle with either water (in the case of L-lactic acid) or mineral oil (all other odorants). The dilutions indicated in the manuscript are the final dilutions considering the fixed air dilution and the dilution of the odorant in the liquid phase.

In most experiments, a recording trial was 10s long, with odor delivered during the 2-3s interval. To check the delivery profile of the odors, a photoionization detector (miniPID 200B, Aurora Scientific) was used. In experiments on the temporal patterns of CO_2_, the trial duration was kept 20 s, and the odor was given in three different temporal patterns: (a) a single pulse of 4s in the 2-6s interval, (b) 4 pulses of 1s each starting at 2s time-point with an inter-pulse interval of 1 s, and (c) 8 pulses of 0.5s each starting at the 2s time-point with an inter-pulse interval of 0.5 s.

### Histology and immunostaining

After electrophysiological recording, the brain was dissected and incubated in 4% paraformaldehyde (Merck, 818715) for a minimum of 30 minutes at room temperature. It was rinsed in phosphate buffer saline (PBS) and incubated in 0.2% of Triton x-100 (Sigma, T8787) diluted in phosphate buffer saline (PBST) for 20 minutes, and then in 5% normal goat serum for 24 hours at 4 degrees or 2-3 hours at room temperature. The primary antibodies used were 1:30 rat anti-N-cadherin (Developmental Studies Hybridoma Bank, DN-EX #8,) and 1:200 rabbit anti-lucifer yellow (Molecular Probes, A-5750). The secondary antibodies used were 1:500 goat anti-rabbit with Alexa fluor 633 (Molecular Probes, A-21070), 1:500 goat anti-rabbit with Alexa fluor 488 (Molecular Probes, A-11008), 1:500 goat anti-rat 405 (Abcam, ab175671), and 1:10^5^ streptavidin with Alexa fluor 488 (Molecular Probes, S-11223) or streptavidin with Alexa fluor 568 (Molecular Probes, S-11226). The brain was washed multiple times in PBS and then bridge-mounted on a glass slide in an anti-fade agent (Vectashield, either pure or diluted in the ratio of 1:1 in 70% glycerol). The mounted brains were imaged using a Zeiss LSM 780 or a Nikon A1R MP+ microscope. All the acquired images are processed using Fiji (ImageJ) software. Glomeruli identification was done manually by comparing images with the antennal lobe atlas for *A. aegypti* provided by Ignell et al. (Ignell et al., 2005), as described previously (Singh et al., 2022).

### Single-sensillum recordings

Single-sensillum recordings were performed in the capitate peg sensilla on the maxillary palps as previously described (Singh et al., 2022). Briefly, an adult female mosquito was immobilized using wax and epoxy resin. A glass recording electrode with 1 µm tip was inserted in a capitate peg sensillum. A reference electrode with 3-4 µm tip was inserted in the eye. Signals were amplified using a Multiclamp 700B amplifier (Molecular Devices) and recorded using a Digidata 1550A digitizer (Molecular Devices).

### Analysis of olfactory responses

All analyses were conducted, and plots were generated using custom code written in MATLAB (Mathworks). We calculated the odor-evoked change in firing rate of a neuron by subtracting the background firing rate (in 0-2s interval) from the firing rate in the response interval (2.1-4.1 s). A duration of 2s was used for the response interval as the odor response to a 1-s odor pulse often lasted more than 1 s. Although the odor pulse was initiated at the 2-s time-point, there was a delay of about 100 ms before the odor reached the animal, and hence we set the response interval to start from 2.1 s. Each odor was typically given for at least 7 trials; the firing rates were averaged over all trials for a cell-odor pair. The odor-evoked change in the membrane potential was calculated in a similar manner using the 40-Hz low-pass filtered recording trace by subtracting the average value between in the 0-2s background interval from the 2.1-4.1s response interval.

To statistically determine whether a cell was responding to an odor, we used the following approach. We divided the response interval into two bins of 1s each and analyzed each bin separately. We created a vector of the firing rate (or the average membrane potential) in a bin over the 7 trials with the odor. Similarly, we created a vector of the responses in the background period over the 7 trials. We then performed a Wilcoxon signed-rank test to check if the responses in the bin were significantly different from the background. When displaying the odor responses of individual PNs or LNs, we used the following graphical notation: a thin horizontal rectangle with two bins is shown at a y-value corresponding to the average firing rate (or the membrane potential), while the color of each bin indicates the statistical significance. If the response in a bin is significantly higher than the background, it is denoted in green color; if lower it is denoted in red; and if there is no difference it is denoted in grey.

In the experiments with temporal patterns of CO_2_, the odor was delivered for a total of 4s in each pattern. The odor-evoked firing rate was calculated by subtracting the average background firing rate from the average firing rate in the 2-10s response interval for each of the three patterns. Firing rates were averaged over the trials.

### Visualization of response trajectories

This analysis was performed to visualize the responses of the whole PN population to different odors. From the 10s recordings, we used the first 6s (including 2s of background, 1s of odor stimulus, and 3s of post-odor period) to visualize the population activity. This period was divided into 60 bins of 100 ms each, with each bin containing the average spiking rate in that bin over the 7 trials with an odor. From a population of 151 PNs, we obtained a 151×60 matrix of responses for the odor, which tells how the 151-dimensional population response evolves over the 60 bins. For easier visualization, the 151×60 matrix is compressed into a 3×60 matrix using principal component analysis. The reduced 3-dimensional responses over the 60 time-points were connected to plot a response trajectory for each odor.

### Classification analysis

The purpose of this analysis was to check if the neuronal responses to the mixture were different from the responses to the two individual odors (CO_2_ and L-lactic acid). The response interval of 2s was divided into two 1s bins, and the firing rate in each bin (minus the background firing rate) was calculated for each of the 7 trials, thus giving a 7×2 matrix for each cell-odor pair.

We first performed the classification analysis for individual PNs. We iteratively used one of the 7 trials as the test trial and the remaining 6 trials to create a training template for each odor by taking the average of the responses over the 6 trials. Next, we calculated the Euclidean distances between the mixture response in the test trial and the three templates. If the mixture response in the test trial was closest to the mixture template, the accuracy in that trial was taken as 1, otherwise as 0. This procedure was repeated 7 times with different test trials, and accuracy values were averaged to obtain the final accuracy value for the PN. Note that the chance level (i.e., expected accuracy for random data) is 1/3 as 3 templates are compared every time. A classification accuracy value significantly greater than 1/3 indicates that the mixture responses across trials are consistently different from the responses to the individual odor components.

To perform this analysis using PN population responses, we concatenated the 2-bin long response vectors from all 151 PNs, separately for each of the 7 trials, thus giving a 7×302 matrix for each odor. After this, we performed the same procedure as mentioned above, by using the 302-long response vectors in the Euclidean distance calculation. The accuracy values were calculated iteratively by using each of the 7 trials as the test trial.

### Prediction accuracy for the temporal patterns of CO_2_

This analysis was similar to the classification analysis described above but performed with the 3 temporal patterns (1 pulse, 4 pulses, 8 pulses) of CO_2_ stimulus instead of three different odors. The analysis was performed separately for each of the 9 PNs in which the three stimulus patterns were tested. For each stimulus pattern, we calculated a binned response vector of length 8 containing the firing rates (minus the background firing rate) in every 1s bin in the 8s response period (2.1-10.1s interval in the trial). As each stimulus pattern was tested for 7 trials, we obtained a matrix of size 7×8 for each of the three stimuli for a given PN. Then, by using one trial as the test trial and the remaining trials to create templates, we checked the distances of each of the test response to the three templates. If a test response (from one of the stimulus patterns) was closest to its corresponding template, the accuracy was taken as 1, otherwise as 0. By iterating over the 3 response patterns and then over the 7 trials as the test responses, we obtained 21 accuracy values, which were then averaged to obtain a final accuracy value per neuron.

### Statistics

Comparisons between samples were performed using the non-parametric Wilcoxon signed rank tests. The tests were two-tailed and were implemented using the *signrank* function in MATLAB (Mathworks) with default parameters.

## Acknowledgements

We thank Arun Shankar for help with breeding mosquitoes and Kazumichi Shimizu for advice on electrophysiology and histology. We thank NG lab members for feedback on the manuscript. This work was supported by the DBT/wellcome Trust India Alliance Fellowship [grant number IA/I/15/2/502091], DST/SERB Swarnajayanti Fellowship [SB/SJF/2021-22/04-C], and SERB Core Research Grant [CRG/2020/004719] awarded to NG.

## Authors Contributions

Conceptualization: NG, Shefali G

Methodology: Shefali G, NG

Data collection: Shefali G, PS, SSS,

AKG Data curation: Shefali G, PS, SSS

Data analysis: Shefali G, MG,

Smith G, SSS, PS,

NG Writing: Shefali G, NG

