## Supplementary Information for "Representations of carbon dioxide in the mosquito antennal lobe"

Goyal et al.

### Supplementary Figure S1

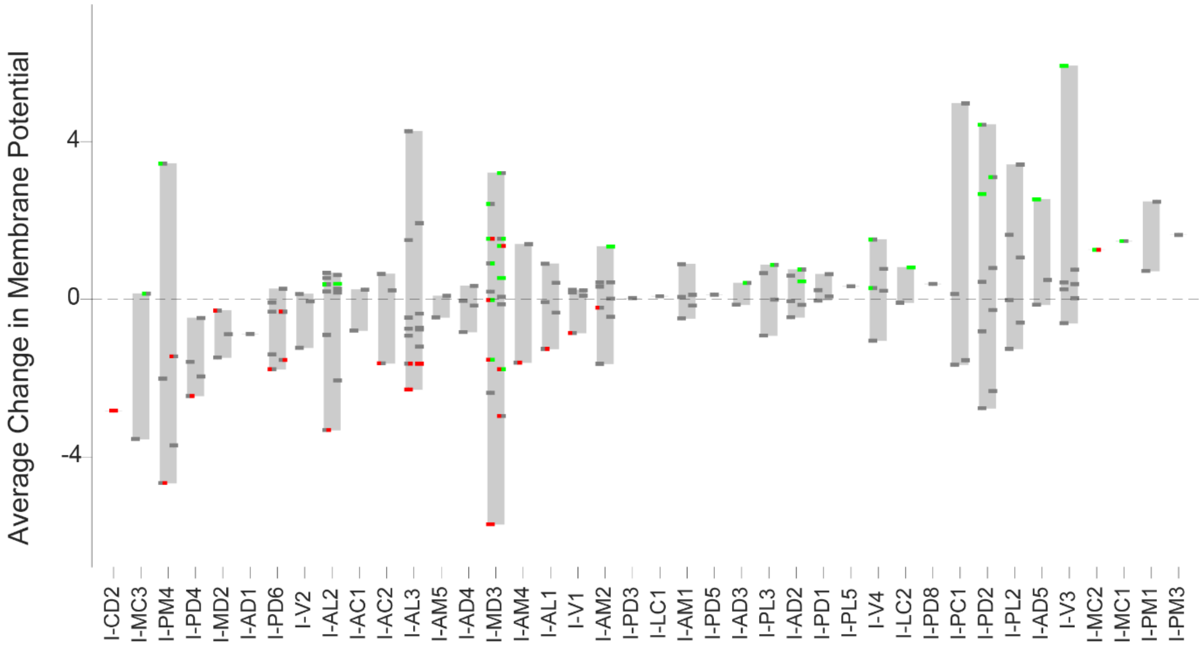


**Supplementary Figure S1: Membrane potential responses of PNs to $CO_2$**

Changes in the membrane potential of PNs, organized according to their glomerular identity, in response to a 1s pulse of $CO_2$. The graph uses the same set of PNs and the same representation as used in Figure 1a.

### Supplementary Figure S2

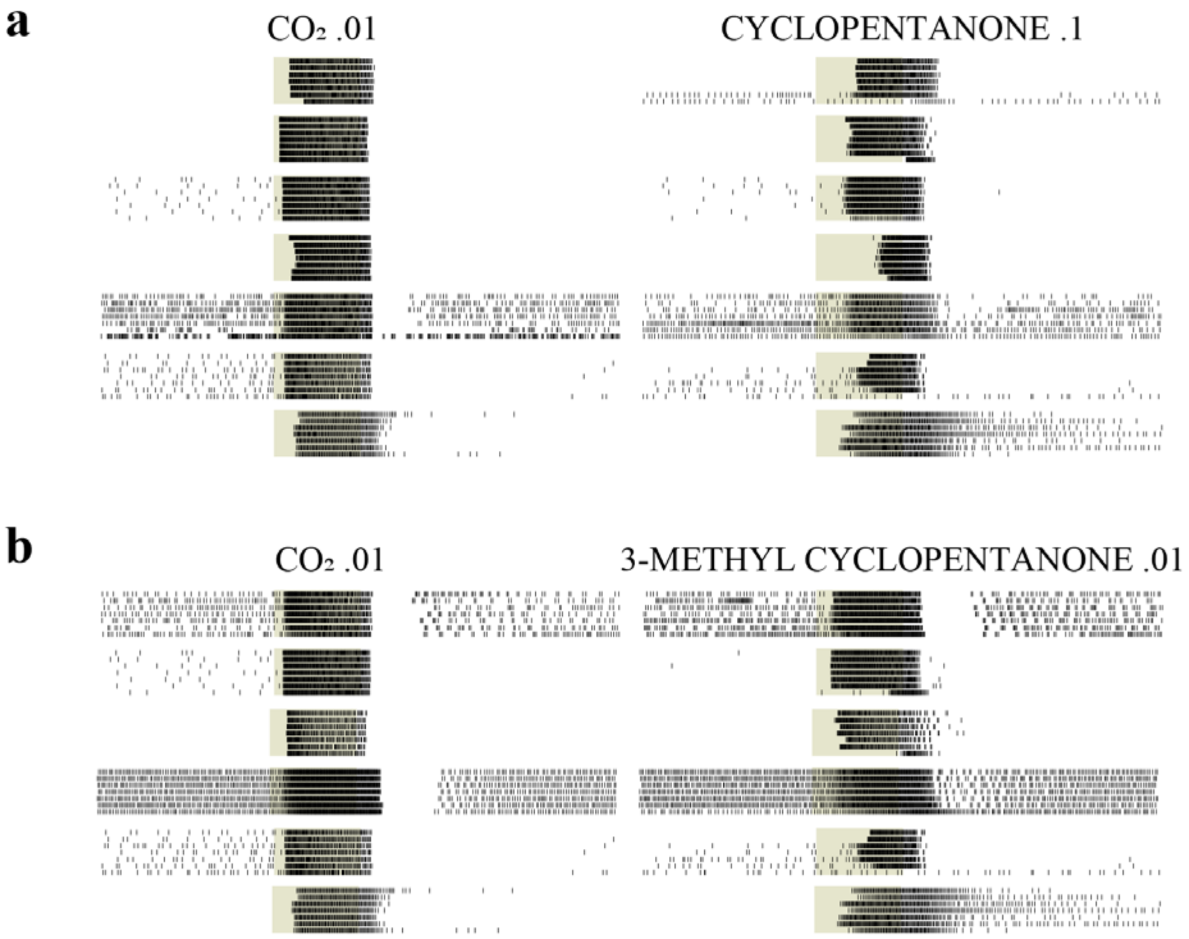


**Supplementary Figure S2: Comparison of cpA responses to $CO_2$ with responses to cyclopentanone and to 3-methyl cyclopentanone**

**a** A side-by-side comparison of the spiking responses of cpA neurons, measured using single sensillum recordings from the capitate peg sensilla on the maxillary palps, to 1s pulses of $CO_2$ and cyclopentanone 0.1. The responses in the same row are from the same sensillum. **b** A similar comparison between cpA responses to $CO_2$ and 3-methyl cyclopentanone 0.01. Some sensilla are common between **a** and **b**. Odor timing is indicated by the shaded background.

### Supplementary Figure S3

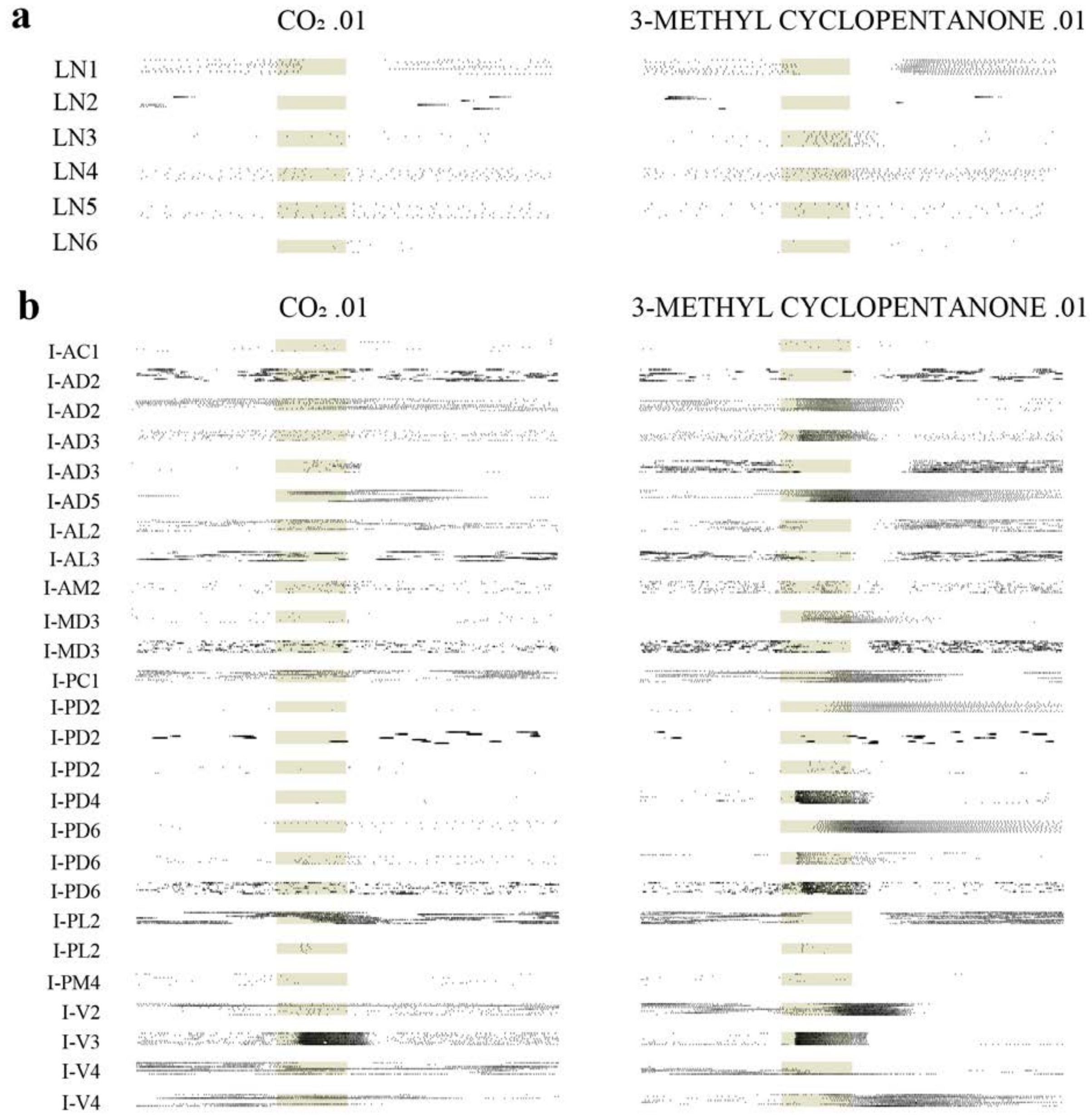


**Supplementary Figure S3: Responses of antennal lobe neurons to $CO_2$ and 3-methyl cyclopentanone**

A side-by-side comparison of the spiking responses of LNs (**a**) and PNs (**b**) to 1s pulses of $CO_2$ and cyclopentanone 0.1. Odor timing is indicated by the shaded background.

### Supplementary Figure S4

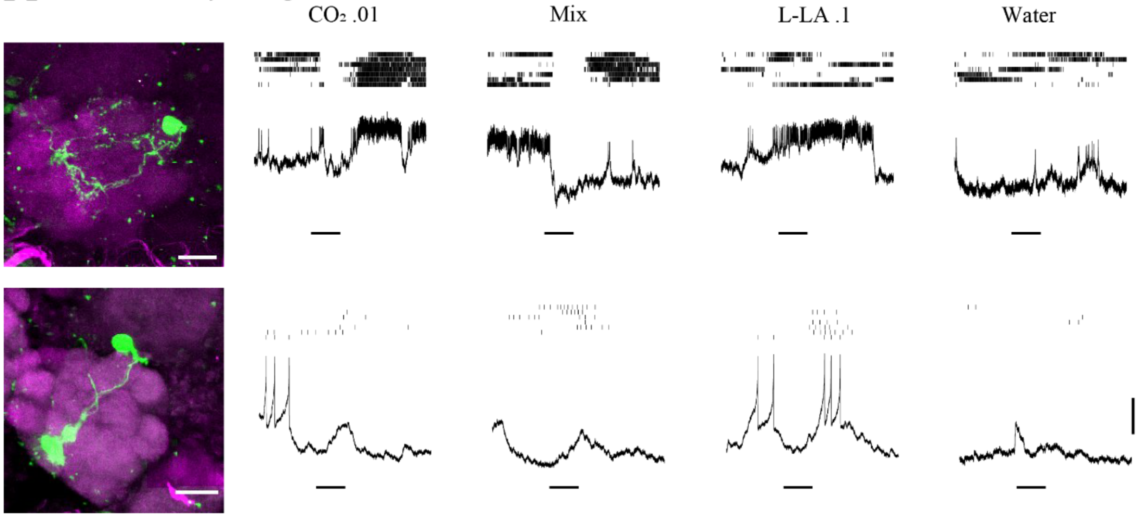


**Supplementary Figure S4: Responses of a multiglomerular and a biglomerular PNs to $CO_2$, L-lactic acid, and the mixture**

Morphologies (within the antennal lobe) and odor responses of a multiglomerular PN (top) and a biglomerular PN (bottom) to 1s pulses of $CO_2$, L-lactic acid (L-LA), their mixture (mix) and water (solvent control for L-lactic acid). Scale bar in morphological images, 20 µm.
